# The Sheep and the Goats: Distinguishing transcriptional enhancers in a complex chromatin landscape

**DOI:** 10.1101/324582

**Authors:** Anne Sonnenschein, Ian Dworkin, David N. Arnosti

## Abstract

Predicting regulatory function of non-coding DNA using genomic information remains a major goal in genomics, and an important step in interpreting the cis-regulatory code. Regulatory capacity can be partially inferred from transcription factor occupancy, histone modifications, motif enrichment, and evolutionary conservation. However, combinations of these features in well-studied systems such as *Drosophila* have limited predictive accuracy. Here we examine the current limits of computational enhancer prediction by applying machine-learning methods to an extensive set of genomic features, validating predictions with the Fly Enhancer Resource, which characterized the transcriptional activity of approximately fifteen percent of the genome. Supervised machine learning trained on a range of genomic features identify active elements with a high degree of accuracy, but are less successful at distinguishing tissue-specific expression patterns. Consistent with previous observations of their widespread genomic interactions, many transcription factors were associated with enhancers not known to be direct functional targets. Interestingly, no single factor was necessary for enhancer identification, although binding by the ′pioneer′ transcription factor Zelda was the most predictive feature for enhancer activity. Using an increasing number of predictive features improved classification with diminishing returns. Thus, additional single-timepoint ChIP data may have only marginal utility for discerning true regulatory regions. On the other hand, spatially- and temporally-differentiated genomic features may provide more power for this type of computational enhancer identification. Inclusion of new types of information distinct from current chromatin-immunoprecipitation data may enable more precise identification of enhancers, and further insight into the features that distinguish their biological functions.

## INTRODUCTION

Enhancers, cis-regulatory elements that coordinate the input of transcriptional activators and repressors, are the primary determinants of eukaryotic gene expression. Enhancers can regulate genes from distances ranging from a hundred base pairs to hundreds-of-thousands of base pairs, and genes are frequently regulated by multiple enhancers to provide complex spatial and temporal regulation (Small et al. 1993, Yao et al. 2008, Pennacchio et al. 2013). Changes in enhancers play pivotal roles in the evolution of multi-cellular life (Carroll 2008, Wittkopp and Kalay 2012, Shlyueva et al. 2014), and disruption in enhancer function has been linked in many cases to disease (Emilsson 2008, Epstein 2009, Corradin et al. 2013, Pennachio et al. 2013).

Identifying the complete repertoire of enhancers for a specific gene is difficult, although there are a number of features that can provide insight. Enhancers are frequently characterized by enrichment of binding motifs for various transcription factors (Halfon et al. 2002, Stathopoulos et al. 2002, Narlikar and Ovcharenko 2009). This may be accomplished using prior knowledge of binding affinities for specific factors and curated position weight matrices (PWMs) (Bailey and Elkan 1995, Stormo 2000), or by searching for over-represented motif clusters in sequences adjacent to co-expressed genes (Sinha et al. 2004, Elemento et al. 2007). However the presence of motifs does not indicate when an enhancer would be active in development, and motifs are not a perfect indicator of transcription factor binding (Junion et al. 2012). Genome-wide studies have used features such as chromatin immunoprecipitation data for transcription factor occupancy and histone modifications to infer function of non-coding DNA; however, features associated with enhancers are also found in regions of open chromatin that are not necessarily active (Fisher et al. 2012, Graur et al. 2013, Spivakov 2014). Many enhancers will have features of regulatory DNA even while inactive (Ghavi-Helm et al. 2014) and transcription factors and polymerases may bind in preparation for future activity (Kleftogiannis et al. 2015b).

Going beyond simple predictions of overall activity, several studies have attempted to predict spatial and temporal aspects of enhancer function. These have had some success at classifying enhancers into broad categories (Zinzen et al. 2009, Wilczynski et al. 2012, Cannavo et al. 2015), and making inferences about the frequency and distribution of specific types of enhancers (e.g. ′shadow enhancers′) (Cannavo et al. 2015). However, results from thermodynamic models used to predict specific expression patterns based on transcription factor binding have shown that achieving that level of specificity is a challenging problem (Dresch and Arnosti 2015, Sayal et al. 2016).

After identifying a prospective enhancer, an additional challenge is determining which gene or genes it regulates (Kleftogiannis et al. 2015b). Enhancers can be very distal to their target gene, which may not be the closest transcriptional unit,(Frankel et al. 2010, Montavon et al. 2013, Claussnitzer et al. 2015) and a single enhancer can influence multiple genes (Frankel et al. 2010, Perry et al. 2010, Cannavo et al. 2016). Most computational approaches do rely on proximity, as a substantial percentage of enhancers control the nearest gene (Sanyal et al. 2012, Shen et al. 2012, Whitaker et al. 2014). However, since many enhancers direct distal genes, this heuristic is far from universally applicable (Whalen et al. 2016). Chromatin conformation data can provide more direct information about which genes are regulated by enhancers (Ernst and Kellis 2012, Shlyueva et al. 2014, Whitaker et al. 2014). Many of these experiments have been conducted in cell-culture, but increasingly datasets from specific organ systems are also becoming available (Williamson et al. 2014). However, identifying tissue specific enhancers from chromatin conformation data is complicated by the large number of non-transcriptional interactions between genomic regions (Mora et al. 2016). Combinatorial protein binding can also provide insight into enhancer locations; clusters of transcription factor binding can indicate enhancer activity. Genes known to be involved in a developmental network are likely to be associated with enhancers bound by common transcription factors regulating that network (Markstein et al. 2004, Zeitlinger et al. 2007).

In *Drosophila melanogaster* enhancer identification and characterization has traditionally relied on genetic analysis (e.g. Fu et al. 1998) and in-vivo reporter assays (e.g. Small et al. 1993). However, even in organisms with relatively small genomes like *Drosophila*, this is a daunting task on a genome-wide scale (Wang et al. 2013). High-throughput methods that assay the entire genome, such as STARR-seq, offer a more comprehensive perspective, but have thus far been confined to analysis of potential enhancers in cultured cells, and only provide insight on transcriptional activity, not function in control of specific genes. Most of these assays are also limited to showing enhancer readout on specific basal promoters (which can have a substantial impact on enhancer responsiveness), and neglect the influence of genomic neighborhood (Arnold et al.2013, Zabidi et al. 2015, Arnold et al. 2017). Evolutionary conservation has been successfully used to identify *Drosophila* enhancers where functional regions of non-coding DNA are more highly conserved than background (Emberley et al. 2003, Glazov et al. 2005, Marcovitz et al. 2016), although this method is complicated by the relatively uniform conservation across most upstream and intronic regions (Halligan and Keightley 2006). Furthermore, in many cases enhancers have conserved function despite sequences being highly diverged (Hare et al. 2008), and non-coding regions with functions unrelated to enhancer activity can also be conserved above background (Glazov et al. 2005, Zhang et al. 2006, Ahituv et al. 2007).

Ultimately, no one genomic feature is an exclusive or universal indicator of enhancer location or activity (Wang et al. 2013). The increasing availability of genomics datasets for enhancer correlates, like nucleosome occupancy and transcription factor binding (Roy et al. 2010), as well as spatial information from techniques like HiC (Lieberman-Aiden et al. 2009) and ChIA-Pet (Li et al. 2010), has opened up the door for compound approaches wherein multiple features are used to infer the activity of enhancers (e.g. Erwin et al. 2014, Capra 2015). Machine learning approaches which combine multiple genomics datasets have employed a broad range of supervised and unsupervised methods, as well as different combinations of enhancer features (Li et al. 2016). Some studies have also successfully assigned enhancers to genes using a probabilistic approach, where both proximity and biological information are considered (He et al. 2014, Kleftogiannis et al. 2015a). Taken together, these studies suggest that compound approaches, using multiple features and diverse types of features as predictors are the most successful (Griffon et al. 2015, Kleftogiannis et al. 2015a, van Duijvenboden et al. 2016). However, distinguishing between functional binding from background represents a major challenge to using genome-wide datasets to predict enhancer activity, and the accuracy of enhancer predictions is not entirely known. Many efforts at estimating the accuracy of enhancer predictions have been limited to testing a handful of representative cases (e.g. Gurdziel et al. 2016), or correlations in place of broad-scale verification. Enhancers are identified based on correlating features (e.g. motifs, transcription factor occupancy), and the accuracy of these calls are estimated from different correlates (e.g. DNase hypersensitivity, (Fletez-Brant et al. 2013, Griffon et al. 2015) and eRNAs, (Chae et al. 2015). Until recently, more direct methods for estimating accuracy have not been available (Coppola et al. 2016, van Duijvenboden et al. 2016).

The recently published Fly Enhancer Resource is a database of information for nearly 8000 *Drosophila* genomic regions, showing the activity of in-vivo reporters with these regions at various developmental stages. It is also an ideal resource for determining both the predictive power of different features associated with enhancers. Importantly, the database provides detailed information about active enhancers throughout development, as well as many genomic regions that are not transcriptionally active. This data can be used as a training set for distinguishing active enhancers from inactive regions, and determining which features are most effective for distinguishing these groups. Here, we exploit the possibilities for validation provided by the extensive functional survey by Kvon et al. 2014, and use data on transcription factor occupancy, chromatin marks, and DNA sequence to push the limits in enhancer identification and classification, and explore what conditions could best be used to distinguish this state.

## 1 METHODS

### A. Datasets used as predictive features

Predictive features were obtained from a variety of publicly available datasets (see Table 1, formatted files available in File S1). For each dataset, information that overlapped with the coordinates of DNA elements were categorized as features of those DNA elements. These include conservation information between *D. melanogaster* and other species in the Drosophila genus, motif scores based on position weight matrices (PWMs) associated with transcription factors involved in dorsal-ventral embryonic patterning, and chromatin immunopreciptation (ChIP) data for a number of transcription factors and histone modifications. ChIP data all came from stage 4-6 embryos; we chose to focus on early embryonic development, as the regulatory networks governing this stage of development are extremely well understood, and many genes at this stage are expressed in simple, easily categorized expression patterns. This stage also has the fewest active genes and the smallest number of active DNA elements (as indicated in Kvon et al. 2014), which simplifies the task of assigning enhancers to genes.

**Table 1.** Features for enhancer classification

| ID | Short ID | Features | Reference |
| --- | --- | --- | --- |
| Dorsal 2015 ChIP-seq | DI 15 s | 1748 | Sun et al. 2015 |
| Dorsal 2009 ChIP-chip | DI 09 c | 9357 | Macarthur et al. 2009 |
| Snail 2014 ChIP-chip | Sna 14 c | 7735 | Rembold et al. 2014 |
| Snail 2009 ChIP-chip | Sna 09 c | 596, 2800 | Macarthur et al. 2009 |
| Twist 2014 ChIP-chip | Twi 14 c | 8629 | Rembold et al. 2014 |
| Twist 2011 ChIP-seq | Twi 11 s | 8797 | He et al. 2011 |
| Twist 2009 ChIP-chip | Twi 09 c | 6684, 7414 | Macarthur et al. 2009 |
| Bicoid 2013 ChIP-seq | Bcd 13 s | 2061 | Paris et al. 2013 |
| Bicoid 2009 ChIP-chip | Bcd 09 c | 619, 702 | Macarthur et al. 2009 |
| Caudal 2010 ChIP-seq | Cad 10 s | 4045 | Bradley et al. 2010 |
| Caudal 2009 ChIP-chip | Cad 09 c | 1590 | Macarthur et al. 2009 |
| Hunchback 2013 ChIP-seq | Hb 13 s | 4986 | Paris et al. 2013 |
| Hunchback 2009 ChIP-chip | Hb 09 c | 1831, 1717 | Macarthur et al. 2009 |
| Giant 2013 ChIP-seq | Gt 13 s | 4194 | Paris et al. 2013 |
| Giant 2009 ChIP-chip | Gt 09 c | 1069 | Macarthur et al. 2009 |
| Hairy 2009 ChIP-chip | Hry 09 c | 1704, 2729 | Macarthur et al. 2009 |
| Knirps 2010 ChIP-seq | Kni 10 s | 505 | Bradley et al. 2010 |
| Kruppel 2013 ChIP-seq | Kr 13 s | 5309 | Paris et al. 2013 |
| Zelda 2011 ChIP-seq | Zld 11 s | 9432 | Harrison et al. 2011 |
| H3K27ac 2015 ChIP-seq | H3K27ac 15 | 3055 | Kok et al. 2015 |
| H3K27ac 2010 ChIP-seq | H3K27ac 10 | 3658 | Roy 2010 |
| H3K4me1 2015 ChIP-seq | H3K4me1 15 | 1934 | Kok et al. 2015 |
| p300 2010 ChIP-seq | p300 10 | 3296 | Roy et al. 2010 |
| Zelda motif sanger | Zld m1 | 17179 | Fly Factor Survey |
| Zelda motif solexa | Zld m2 | 19271 | Fly Factor Survey |
| Dorsal motif FlyReg | DI m1 | 71087 | Fly Factor Survey |
| Dorsal motif NBT | DI m2 | 26159 | Fly Factor Survey |
| Snail motif FlyReg | Sna m1 | 38221 | Fly Factor Survey |
| Snail motif sanger | Sna m2 | 31059 | Fly Factor Survey |
| Snail motif solexa | Sna m3 | 42341 | Fly Factor Survey |
| Twist motif da | Twi m1 | 29656 | Fly Factor Survey |
| Twist motif FlyReg | Twi m2 | 61676 | Fly Factor Survey |
| UCSC BlastZ scores | NA | NA | Rosenbloom et al. 2015 |

Pairwise alignments were downloaded in axt file format from the University of California Santa-Cruz Genome Browser (Rosenbloom et al. 2015) (http://hgdownload.soe.ucsc.edu/downloads.html). Alignments used were *D. melanogaster* version dm3 to *D. grimshawi* droGri2,*D. willistoni* droWil1, *D. ananassae* droAna3, *D. pseudoobscura* dp4, *D. erecta* droEre2, *D. yakuba* droYak2, *D. sechellia* droSec1, and *D. simulans* droSim1. These were chosen to reflect a range of phylogenetic distances from *D.melanogaster*. Summary lines from each chromosome axt file were used to create a single summary file for each genome (See File S1). A custom python script was used to determine the average BLASTZ score per 100 base pairs for a region, which was plotted by species on a log10 scale. Perfect conservation over 100 base pairs would yield a score near 10,000. Scripts and intermediate files are available through Figshare in Files S1 and S2. BLASTZ scores were used for pairwise comparisons (Visel et al. 2007).

Position probability matrices for the transcription factors Dorsal, Snail, Twist and Zelda were obtained from the database Fly Factor Survey (Zhu et al. 2011). These values were formatted using a custom python script to enable them to be input into the MEME Suite program MAST (Bailey and Gribskov 1998). Motif matches throughout the genome with a p-value of less than 0.0001 were obtained using MAST, and the release 5.37 version of the *Drosophila melanogaster* genome. Output files, containing coordinates of qualifying matches and their scores, were converted to bedgraph format.

To obtain insight into factors driving enhancer activity, we used ChIP-chip and ChIP-seq data. Peak scores were obtained from public databases including ModEncode, the Berkeley *Drosophila* Genome Project, and other publications with relevant ChIP data (see Table 1 in Methods). In each case terminal data files were converted to bedgraphs (available in File S1). When necessary, coordinates were converted to the release 5.37 genome using the Flybase coordinate conversion tool. As these datasets were analyzed using different pipelines, they are somewhat variable regarding distributions of scores, and thresholds for what was considered a significant peak. The overlap (of at least one nucleotide) between peaks called by different datasets, and correlation between scores in overlapping peaks, was analyzed using a custom python script, and visualized using gplots (version 3.0.1) in R (version 3.3.2).

### B. Organization and curation of data from Fly Enhancer Resource

DNA elements from the Stark Lab Fly Enhancer Resource were downloaded from the Kvon et al. 2014 supplementary tables. For comparisons made in stage 4-6 embryos, we used regions that were active in stage 4-6 embryos, filtered to include only those with manually-verified activity scores of 3 or higher, and confirmed inactive regions in stage 4-6 (annotated by Kvon et al. 2014). These are listed in File S3. For examination of expression patterns, the highly active stage 4-6 DNA elements were re-annotated based on photographs of the slides, to identify those that exclusively fit characteristic Anterior-Posterior and Dorsal-Ventral expression patterns. 114 DNA elements were found to have strong expression distinctly in the Anterior region of the embryo, while 78 were only expressed in the Central or Posterior regions. 22 DNA elements were identified that had stereotypical Mesodermal or Neurogenic Ectoderm expression patterns (as in enhancers for *snail* or *short gastrulation* respectively), and 192 had strong expression patterns that did not overlap with these two regions. All active and inactive DNA elements that Kvon et al. 2014 identified as ′verified′ were used for classifications based on temporal specificity (e.g. ubiquitously active, or stage specific). DNA elements were grouped by whether they were active in all embryonic stages, only in stages 4-8, stages 9-12, stages 13-16 (Figure 7).

### C. Intersection of features with Fly Enhancer Resource

To determine if these features could be used to distinguish between spurious binding and functional binding, we compared the distribution of peak scores for features that overlapped (by at least one base pair) with genomic regions that drive strong expression in stage 4-6 embryos (as reported by the Fly Enhancer Resource) and regions that do not drive expression in stage 4-6 embryos. Every DNA element tested by the Fly Enhancer Resource was assigned a score for each feature included. Scores from bedgraph items (peak scores in the case of ChIP datasets, MAST motif scores in the case of motif datasets, and BLASTZ scores for conservation information) that overlapped with a DNA element were assigned to form that DNA element′s scores (multiple peaks within an element were combined; scores were only combined within a single dataset, thus normalization was not necessary between datasets). If a DNA element did not overlap with any features for a given dataset, its score for that feature was set to a numerical value one standard-deviation lower than the lowest score that indicated a hit for that dataset, based on the distribution of values for that feature. We calculated fractional overlap of these genomic features, both to determine whether individual features were reproducibly identified in different studies, and to see if there was overlap of features known to be functionally related. Each dataset was compared pair-wise with every other dataset; the degree of feature overlap was calculated as a percentage (the percentage of the features in the smaller dataset that overlap with one or more features in the larger dataset). In Fig. 10 and Figure S7, the Fly Enhancer Resource DNA elements were filtered to only include those that fit a minimal threshold for a potential active enhancer, based on binding of transcription factors from at least one of the datasets described in Table Subsets of this set of potential enhancers were created to allow comparisons between active DNA elements and inactive regions that resemble active DNA elements with regards to transcription factor occupancy. This was done by restricting the data to include only elements bound by two or more transcription factors, or only DNA elements occupied by three or more transcription factors (see Results). All sets of DNA elements were under-sampled from the larger category to balance the number of active and inactive DNA elements included in these prospective training sets.

The DNA elements with features assigned were scaled and visualized using a Principal Components Analysis using the stats package (version 3.2.3) in R (version 3.3.2). This code is available in File S4. The percentage of variation captured is displayed in a scree plot (Figure 5b).

### D. Discrimination between classes of enhancers with Random Forest

To determine if it is possible to distinguish between regions that are transcriptionally active and regions that are soon-to-be active, we also classified the Fly Enhancer Resource DNA elements into several categories of activity. This included ubiquitous activity (active in every developmental stage measured by Kvon et al. 2014), total inactivity (never active in any measured stage), and stage specific regions that are active only in early, middle, or later embryonic stages. To facilitate comparisons of different sorts of data, values for ChIP, conservation, and motif scores were normalized using the ′scale′ function in R. In Fig. 5, highly active (defined by a score of 3 or higher in stage 4-6) and inactive DNA elements were classified using a random forest algorithm (randomForest package, version 4.6-12) with 500 trees in R (version 3.2.3), using two thirds of DNA elements for training and one third for testing. Receiver Operating Characteristic (ROC), Precision Recall and the area under these curves was analyzed using ROCR package in R (version 1.0-7, Sing et al. 2005). ′Background′, or expected AUC based on random chance, was estimated by creating random values between 1 and 1000 for each feature, and using this as the training set for test set classification. In this and later figures, the DNA elements used for training were randomly re-sampled ten times. In Fig. 6 and Figure S7 where balanced datasets were used equal numbers of active and inactive elements were randomly selected using the package dplyr (version 0.5.0). For subsets filtered for DNA elements containing the indicated ChIP binding protein or chromatin mark, the number of ″active″ elements ranged from 100-300, and for balanced datasets, the same number of inactive elements was used. The same classification was attempted with datasets that were reduced to only include inactive regions that at least superficially resembled active enhancers (regions occupied by two or more transcription factors, or regions occupied by a combination of factors associated with development).

### E. Feature importance analysis

Feature importance was measured using the mean decrease in Gini index within the randomForest function, which estimates variable importance during the training of the random forest. To determine if redundancy or correlation between features influenced importance scores, features were iteratively left out of analyses, so after each set of predictions, the most important or least important feature was dropped. In Fig. 6 where successive features are omitted from the analysis, the most and least important features were re-calculated after each run with subsets of the features. After exclusion of each feature, the order of feature importance frequently changed, as the removal of partially correlated data increased the relative importance of remaining features. Thus, the list of dropped features shown in Table S3 reflects the re-calculated scores for each of the features.

### F. Identifying potential enhancers

In Fig. 10, to comprehensively assess regions that contain bound transcription factors and chromatin signatures associated with enhancers, we generated a list of potential enhancers *de novo* based on clustering of transcription factors adjacent to specific genes of interest, which were selected based on annotated Anterior-Posterior, Mesodermal, or Neurogenic Ectodermal expression patterns. Prospective enhancers were defined by clusters of overlapping features occurring within 50kb windows centered on the +1 position of genes of interest, using a custom python script available in File S2. Genes of interest *brinker (brk), ventral nervous system defective (vnd), short gastrulation (sog), snail (sna), even-skipped (eve), Kruppel (Kr), hunchback (hb)*, and *knirps (kni)*, were selected based on their well studied regulation in early stage embryos. To locate clusters, we defined regions containing at minimum Zelda and one other ChIP signal (p300, H3K4me, H3K27ac, or any TF), or H3K4me1 and one other ChIP signal (as indicated in Fig. 10). The final enhancer was defined as a region around the center of the cluster or clusters of features, and set to a fixed size of 500-2000 bp. These regions were then classified as ′putative enhancers′ or ′background binding′ by random forest algorithms trained on regions from the Fly Enhancer Resource.

### G. Validation of predictions

Annotated enhancers were defined based on verified regulatory regions in the RedFly database (Gallo et al. 2011). Each random forest trained on the Fly Enhancer Resource was used to classify predicted enhancers around genes of interest ten times, using different samples of the training set. A custom python script was used to determine the frequency that each model classified a cluster that overlapped with an annotated enhancer as active (defined as percentage overlap), and the frequency that each model classified non-overlapping segments as active (likely off target). The results of this were graphed with ggplot2.

### H. Data Availability

Supplementary figures, as well as all scripts and files necessary for generating figures, are available on figshare. All genomes features are found in File S1, python code for identifying and assigning features to prospective enhancers is in File S2, annotations from Kvon et al. 2014 and from Redfly are available in File S3, and all R code for visualization and machine learning is in File S4. Scripts and final files are also available on github at https://github.com/asonnens/crm_analysis.

## 2. RESULTS

### A. Enhancer sequences are not highly conserved

Sequence similarity in related species has been extensively used to identify functional non-coding DNA. Highly conserved regions may indicate purifying selection on regulatory elements (Visel et al. 2007), and conversely highly diverged sequences can be a signature of enhancers associated with phenotype differences between species (Gittelman et al. 2015). We looked at sequence conservation between *Drosophila melanogaster* and a range of other species in the Drosophila genus for DNA elements that were either active in every stage of embryonic development or inactive across all stages, to see if there are any visible trends. DNA elements do not show increased sequence conservation over inactive genomic regions as measured by BlastZ scores (Figure S1). This was true for all species comparisons.

### B. Consistency and overlap of distinct chromatin immunoprecipitation data

We collected a range of ChIP datasets to use as indicators of active enhancers. These datasets were produced using different methods (ChIP-chip vs. ChIP-seq) and a range of experimental conditions. Accordingly, the number of features contained within different datasets is highly variable (Table 1). We assessed the consistency and overlap of each dataset to determine how to most effectively combine them. Different datasets measuring binding of the same transcription factor typically did not completely overlap, but did to a greater degree than is seen in comparisons between different transcription factors (Figure 1, Figure S2). Separate datasets produced by the same lab (either at different times, or using different antibodies for the same proteine.g. Twist, Bicoid, and Hunchback) overlapped more than datasets produced by different research groups. MacArthur et al. (2009) reported a 91% overlap in ChIP results using different antibodies for the same transcription factor. When comparing datasets produced by different labs, reproducibility was greater for the highest scoring peaks (one standard deviation above the mean for each dataset). On average, when the same transcription factor was measured by different labs, 57% of the highest scoring peaks were reproducible (Fig. 1), while 52% of all reported peaks (including lower scoring peaks) were in agreement, ranging from 21% overlap between Giant ChIP-chip and ChIP-seq datasets, to 94% of Dorsal ChIP-seq peaks which are also included in the 2009 Dorsal ChIP-chip dataset (Figure S2). Overlap between peaks for different transcription factors ranged from 28% (Kruppel chip-seq with all other datasets) to 56% (Knirps ChIP-seq with all other datasets) averaging at 42 percent. Three out of four Twist datasets and one out of two Snail datasets also showed an unusual degree of cross-correlation with other transcription factors, especially in comparisons of high scoring peaks, as shown in the ′blue stripe′ in Figure 1. Variability between ChIP datasets, even for duplicate measurements of the same factor, has been noted in previous studies (Devailly et al. 2015).

**Figure 1.**
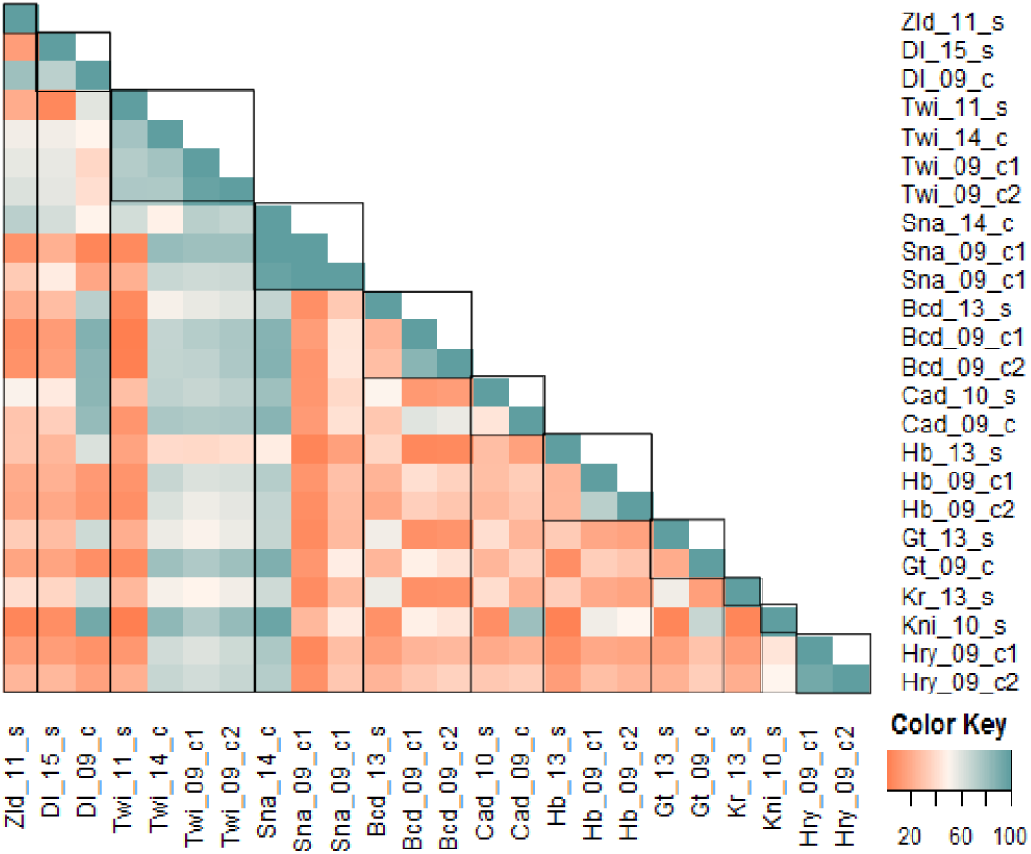
Correlations between genomic regulatory features. Similarities between ChIP datasets are shown by percentage overlap of ChIP peaks between different datasets, ranging from 100 percent overlap (blue) to 0 overlap (red). Boxes on the diagonal highlight independent measurements of specific transcription factors.

### C. Correlations between chromatin immunoprecipitation data and enhancer activity

Although ChIP datasets provide partially inconsistent information about occupancy, in combination they are an important guide to the regulatory potential of specific DNA elements. In some cases, loci that are inactive can be readily distinguished from loci around active genes based on ChIP occupancy alone. Sparse occupancy around weakly transcribed or inactive genes (Figures 2a, 2b) contrasts with abundant binding of factors near loci adjacent to robustly expressed genes (Figure 2c,2d). However, around active genes, some bound regions may not be associated with known enhancers. We found many instances of transcription factor occupancy of tested, inactive regions in the Fly Enhancer Resource (e.g. circled region in 2d). These regions are difficult to visually distinguish from active enhancers.

**Figure 2.**
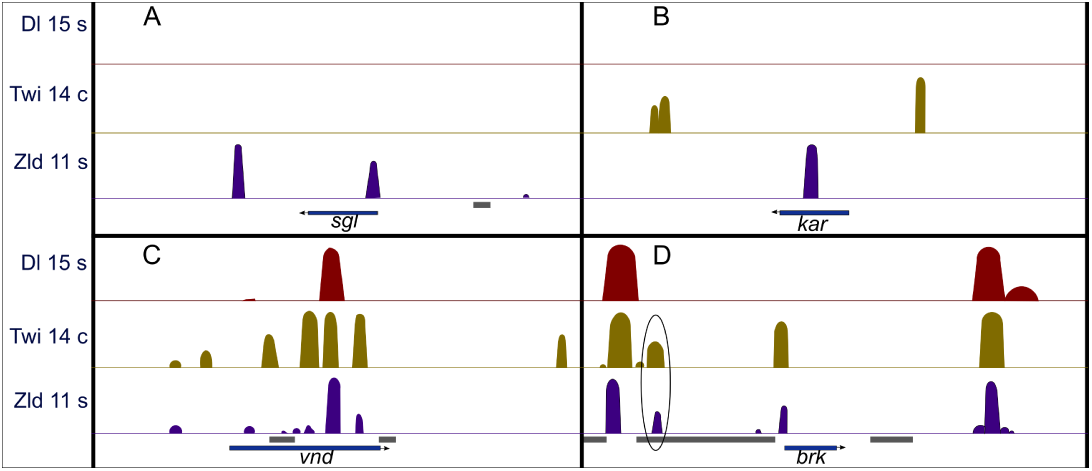
Abundance of ChIP peaks correlates with presence of active regulatory regions, but some clusters do not possess enhancer activity. Low levels of binding by Dorsal, Twist and Zelda at (a) weakly expressed *sgl* locus and (b) inactive *kar* locus.(c) numerous bound regions associated with actively transcribed *vnd* and (d) *brk* loci. However, some regions found to be inactive in Kvon et al. (indicated by underlined genomic regions) are bound by multiple transcription factors (circled region near *brk*).

To determine if ChIP signals that overlapped with active elements had a higher degree of reproducibility, we used the pairwise comparisons of features from Figure 1 to measure the correlation in scores for overlapping features in both DNA elements that were strongly active and inactive in stage 4-6 embryos. In the case of Zelda and Dorsal (transcription factors associated with a large number of ChIP-peaks, Table 1) and histone marks associated with enhancer status, occupancy scores for different datasets measuring the same transcription factors within a given DNA element generally correlated with each other, as did datasets measuring different but functionally related features, although the difference in this correlation between active and inactive regions is minimal (Figure 3). The same trend was largely true for other transcription factors (Figure S3). This suggested that the intensity of peak scores are fairly reproducible between datasets, but that functional binding is not more reproducible or consistent than background binding. Motif scores did not correlate at all with occupancy scores for transcription factors that overlap with the motifs, except in the case of Zelda (Figure 3, Figure S3). This correlation seemed to be much stronger in active DNA elements than in inactive elements (Figure 3c and 3d).

**Figure 3.**
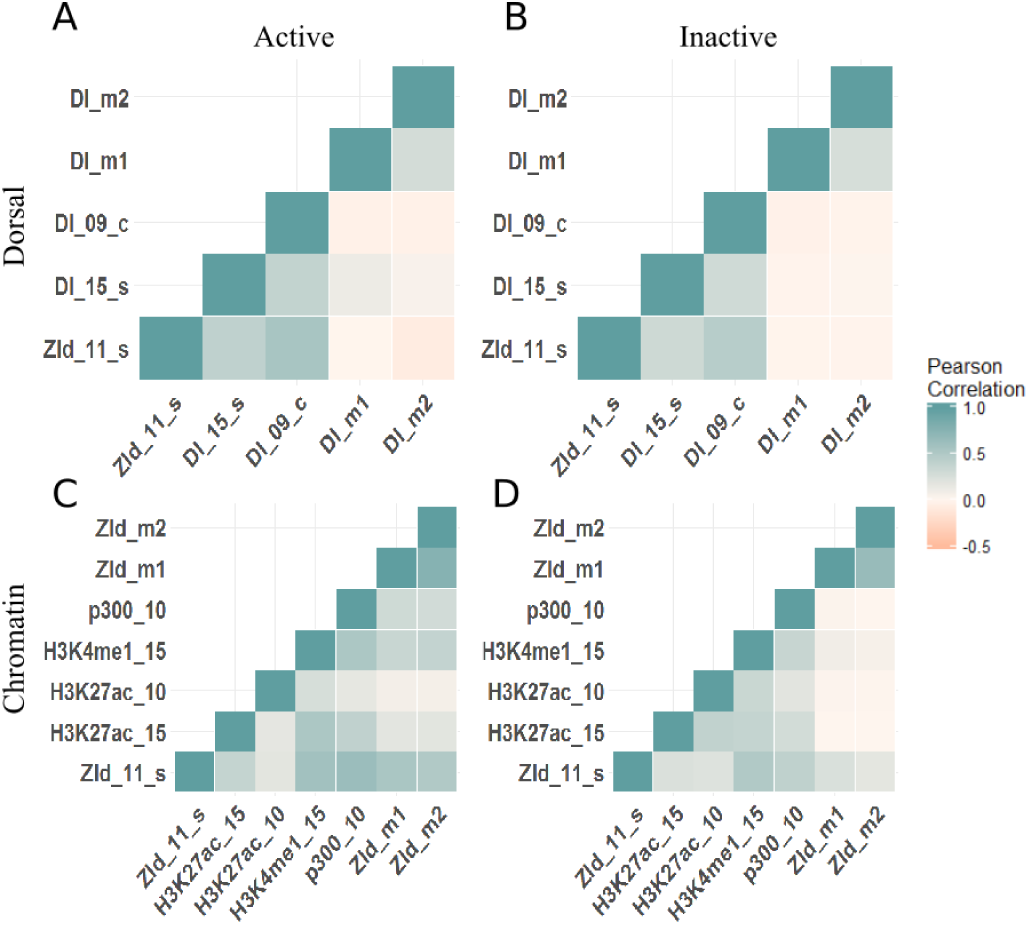
Similar levels of correlation between features for active and inactive elements. (a, b) Levels of Dorsal and Zelda were assessed on active and inactive regions. Zelda and Dorsal peaks were correlated on both classes of elements, but little correlation is noted between protein occupancy and the respective binding motifs. (c, d) For chromatin marks and pioneer factor Zelda protein and motifs, higher correlations were noted on active vs. inactive elements. Results from 392 highly active elements (left) and 6,858 stage 4-6 inactive elements (right). Comparisons for additional factors shown in Figure S3.

The qualitative differences between the ChIP signals around active and inactive elements shown in Figure 2 were less salient when ChIP scores were plotted as a distribution (Figure 4). For some ChIP datasets, the higher-scoring peaks were associated with active DNA elements (e.g. Snail 2014 chip). However this trend was only true for some of the ChIP datasets, suggesting that different thresholds or levels of background binding in these datasets may obscure relevant signals for active vs. inactive binding. No differential signals were apparent for most of the ChIP datasets or for individual binding motifs. DNA elements from both categories that did not overlap with the signal were excluded from the comparison, and the histograms from both categories were scaled to integrate to 1. In most cases, there were a larger number of inactive regions bound by transcription factors than active regions, as there are 6,858 inactive regions vs. 392 active regions (Table S1). Distributions for remaining features are shown in Figure S4.

**Figure 4.**
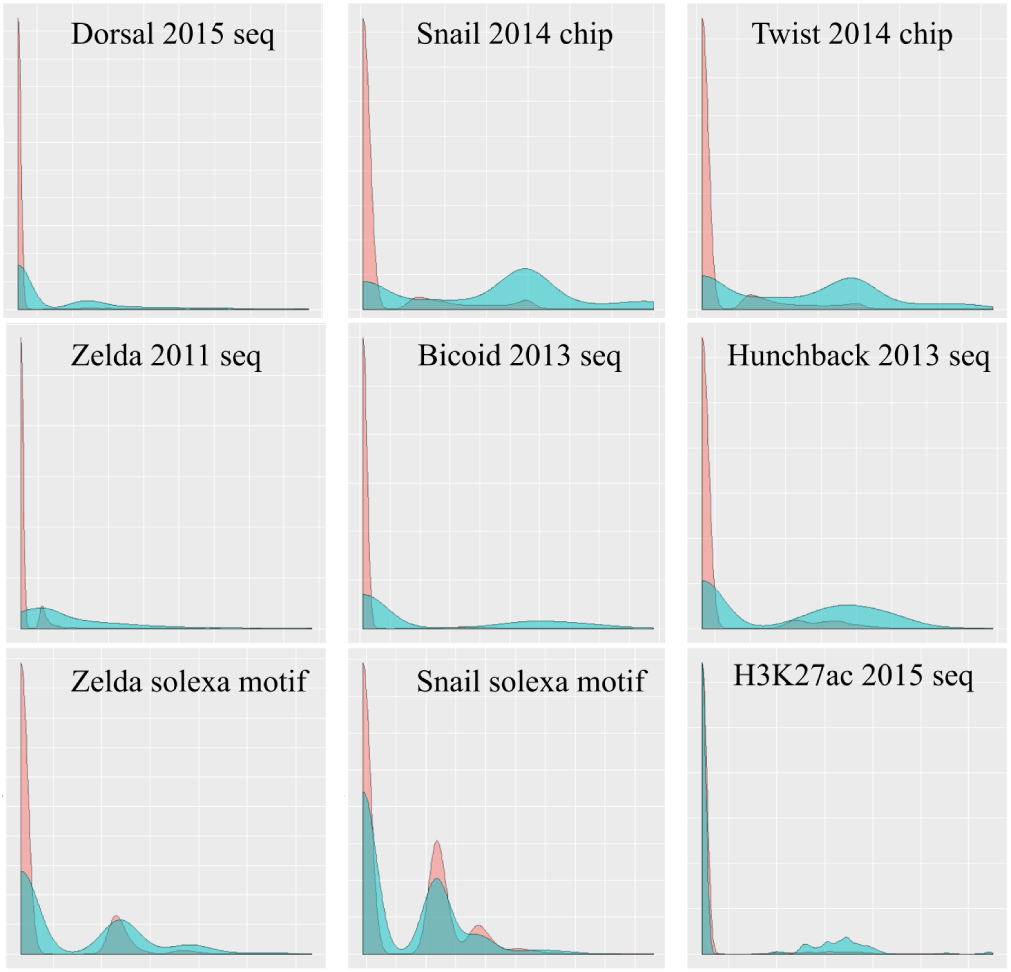
Differential ChIP peak and binding motif enrichment values on active and inactive DNA elements. Histograms showing distributions of ChIP peaks and binding motifs were plotted for DNA elements active (blue) and inactive (red) in stage 4-6 embryos. DNA elements from both categories that did not overlap with the signal were excluded from the comparison. The X axis shows peak or motif score of the relevant protein or histone modification, and Y axis shows frequency scaled to integrate to 1. Distributions for remaining features are shown in Figure S4

### D. Transcriptional activity and stage specificity can be classified based on genomic features

Our analysis indicated that no single feature conclusively correlates with regulatory activity, therefore, we asked if combinations of these features might distinguish active from inactive DNA elements. First to determine the variability of the dataset, we implemented a Principal Components Analysis (PCA) of the forty-one datasets. PCA of all features for all DNA elements (392 active and 6858 inactive) showed some differences in their distribution. (Figure 5a). The largest contributors to PC1 (the greatest sources of variation in the dataset) are mostly occupancy information for transcription factors associated with Anterior-Posterior patterning, like Caudal and Hunchback (Table S2). Interestingly, the largest contributors to PC2 are primarily the degree of evolutionary conservation in each DNA element, based on various species comparisons (Table S2). Almost all variation in the dataset was captured in the first ten principal components (Figure 5b).

**Figure 5.**
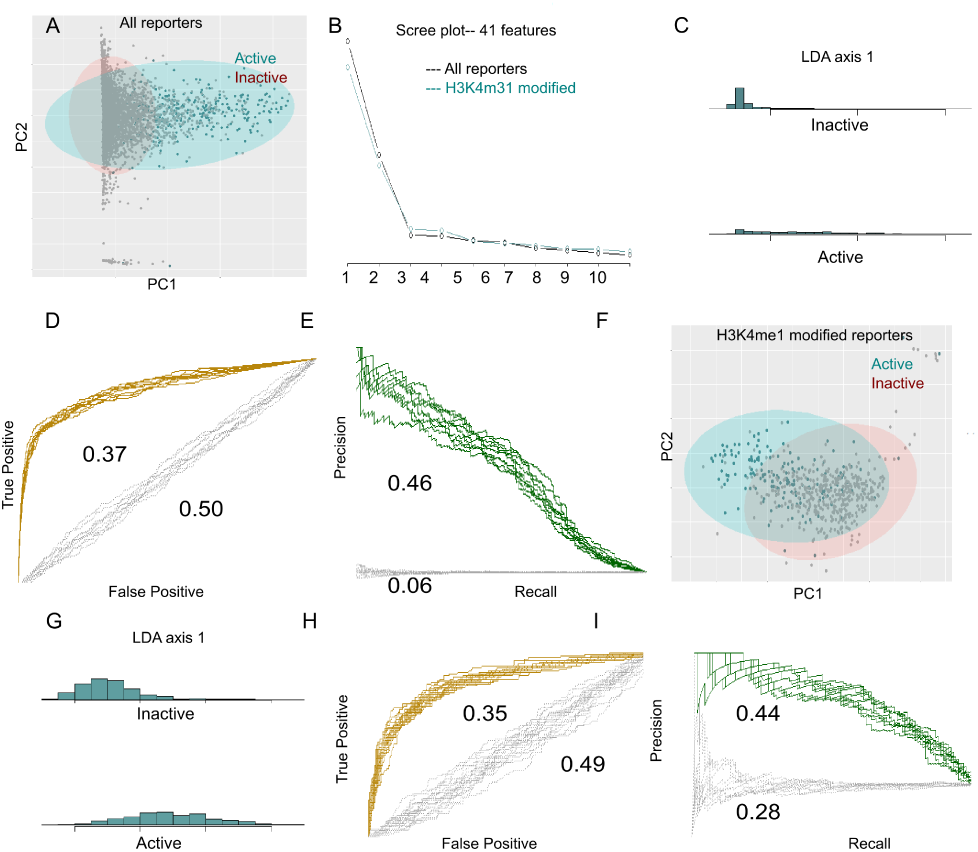
Active DNA elements can be separated from in-active regions. (a) DNA elements with and without regulatory activity can be partially distinguished based on Principal Component 1; additional principal components showed little separation (see Table S2 for loadings and variance for the first six principal components). There is a sharp boundary on the PC1, due to PC1 variation primarily coming from ChIP occupancy scores, and the majority of DNA elements having no occupancy, and thus share an identical ′low score′ numerical stand-in. The separation is far more visible using supervised machine learning. (b) Scree plot for PCA in (a: black) and (f: blue). (c) Separation between active and inactive regions with LDA, using all DNA elements (cross validation in Figure S5). (d) Random forest trained on 2/3 of both unrestricted and restricted datasets (using all 41 features) achieved 96% accuracy of total calls, and a high ratio of true positives relative to false positives, as shown in ROC plots (yellow), and a difference of 0.37 in the area under the curve relative to classifications made on random data with the same sample size (whose ROC are shown in gray). However, Precision Recall (e, green plots) was much lower, although even higher relative to background (gray plots), with a relative AUC difference of 0.46. (f) Using elements that are distinguished by H3K4 mono-methylation reduces the sample size from 392 highly active regions and 6858 inactive regions, to 167 highly active and 421 inactive regions. PCA of these elements showed much less separation between active and inactive. This also lowered the degree of visible separation based on LDA (f) and decreased random forest classification accuracy to 83% and area under the ROC curve (g). It led to substantially higher Precision Recall (i). However, it led to equal increases in the AUC of predictions made on random data of the same sample size, shown in gray.

Although active and inactive regions overlap, the visible partial separation based on the first principal component suggested it may be possible to classify activity using machine learning. Linear discriminant analysis (LDA) of all DNA elements showed a degree of separation, although there was overlap in the distribution of where they fell on LD axis 1 (Figure 5c, cross validation in Figure S5). Next, we used a random forest (chosen for its amenability to biological interpretation) to classify all active and inactive DNA elements based on the same 41 features used for PCA and LDA. A random forest trained on two thirds of the active and inactive DNA elements classified the remaining one third with 96 percent accuracy (measured by total percentage of true positives and true negatives) over 10 trials. Receiver Operating Characteristic (ROC) and Precision Recall curves are shown in Figure 5 d,e. Classifications are much more successful as measured by ROC than by Precision-recall, as indicated by their respective measurements of ′area under curve′ (AUC), although Precision Recall is more improved relative to classifications based on random chance (shown in gray). Zelda and Bicoid datasets were found to be the most important features for classification (Figure S6).

Many of the inactive regions used in the PCA and classifications in Figure 5c-e are relatively sparse in terms of transcription factor occupancy, like the regions shown in Figures 2 a, b. These unoccupied regions can be trivially distinguished from potential enhancers. A more challenging question is how to separate active regions from inactive regions that superficially resemble them, containing many of the same marks, as shown in Figures 2d. Chromatin signatures are frequently used as a proxy for active elements, therefore we selected loci that are associated with H3K4 monomethylation for training and testing, reducing the datasets to 167 highly active regions and 421 inactive regions. These are much less clearly separated by Principal Components (Figure 5f) and LDA (Fig. 5g). Turning to random forests, the AUC for ROC is nearly as high as before (compare Fig. 5d, h), and Precision Recall is dramatically improved (compare Fig. 5e, i). However, the reduced dataset is much closer to having a balanced number of samples for active and inactive elements, so the probability of correct classification due to random chance was greatly increased. Neither ROC nor Precision Recall showed substantial improvements in AUC relative to random chance. Restricting the analysis to loci that were bound by one or more transcription factors similarly improved overall Precision Recall, but not relative to background (discussed in Figure S7).

### E. Impact of number of features on predictive accuracy

To determine if a specific combination of features would be the most effective for predictions (e.g. only using the most predictive features, or excluding the least predictive features), we re-performed random forest classification using the same training data, with different combinations of the 41 predictors. We iteratively left out individual features (Table S3; Figure 6). After each run of ten random forests, the features were ranked using Gini Impurity Index. The next run then excluded either the most or least important feature based on the previous run, and performance was compared between each. We found that dropping the most important feature, or even top three features had little impact on Precision Recall; loss of additional features had some negative impact on classifications. After dropping the eighteen most important features (largely ChIP data), Precision Recall and ROC curves are depressed. However, dropping the least important features had no noticeable impact, until over thirty features are dropped. Analyses using only the top twelve most useful features were equivalent to employing all features. These results suggest that the features must contain a significant amount of redundancy, but the redundancy is not interfering with accurate predictions.

**Figure 6.**
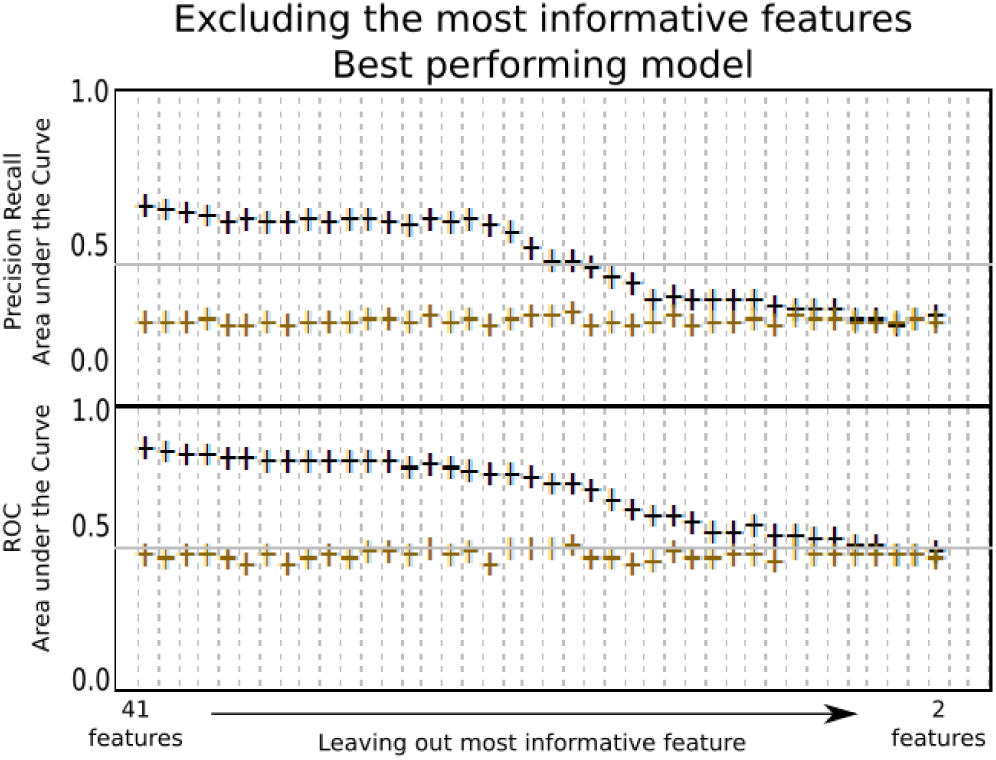
Random forest performance with least and most informative features. Predictive success of models with differing numbers of features, with unbalanced (black) datasets and all active and inactive elements, and balanced datasets (blue), in which 200 active and inactive elements are used for training, and 100 for testing. Predictions were made by iteratively dropping the most informative feature to least informative feature (+), or least informative to most informative (-). These features are listed in order in Table S3. The scores represent predictions made using 41 features, to predictions made using two features. The furthermost right scores are predictions made based on 41 randomly generated features. Note that balanced datasets have substantially higher ′background′, or scores based on random datasets of the same size (as in Fig. 5)

### F. Classifications by stage specificity and expression pattern

Some studies have shown that genes that are active throughout development have features that make them distinct from genes and regulatory regions that are tissue or stage specific (Liu et al. 2017), and that there are additional distinctions between poised enhancers and those that are actively expressed (Creyghton et al. 2010). However, a comparison of DNA elements that are active in all stages with those that are specific to early embryonic development do not suggest that they can be as easily distinguished as active and inactive elements (Figure 8a,c). A possible mitigating factor is that regulatory regions often gain binding of transcription factors and chromatin marks in a successive fashion during development, prior to becoming active. Thus, some of the elements considered above as ″inactive″ may actually be false negatives. Here, some DNA elements are exclusively active in embryonic stages 4-8, whereas others are not active in this earlier period, but become active in stages 9-12 or 13-16 (Figure 7). We tested if the 41 features would distinguish such elements that become active only in later stages based on their features measured at an earlier time point. PCA and LDA analysis did not show much separation (Figures 8b,d), unlike the classification shown in Figure 5. Consistent with these results, random forest classification was only slightly more successful than would be expected by chance (data not shown). The low accuracy may stem in part from the relatively smaller training set, however, it is also possible that the considered genomic features are simply not informative about early vs. late activity in DNA elements. Moreover, most of the features used for this classification were from datasets collected at the early embryonic stage of development; additional information about chromatin conformation or later transcription factor occupancy would likely improve predictions for later enhancers.

**Figure 7.**
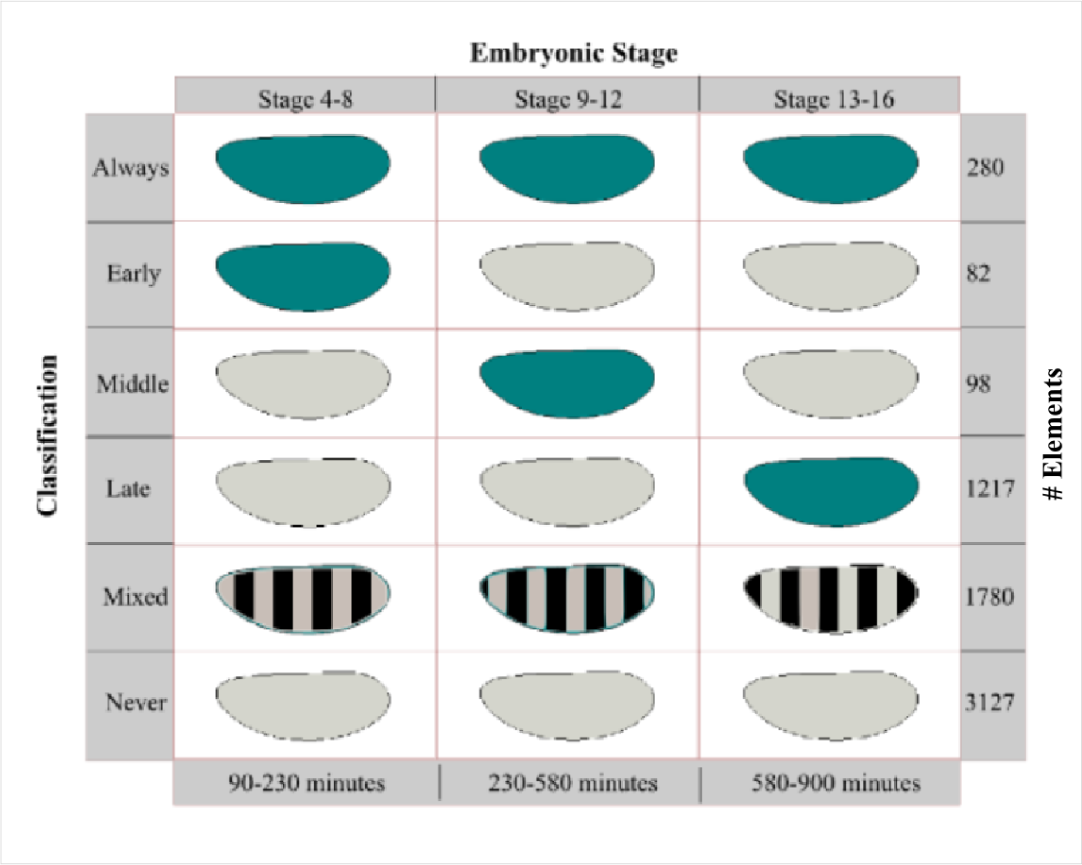
Distinct classes of elements tested for function in embryos. To assess predictions of different classes of active elements, DNA elements (Kvon et al. 2014) were assigned to one of six classifications. ″Mixed″ refers to elements that exhibit activity in two of the three indicated stages.

**Figure 8.**
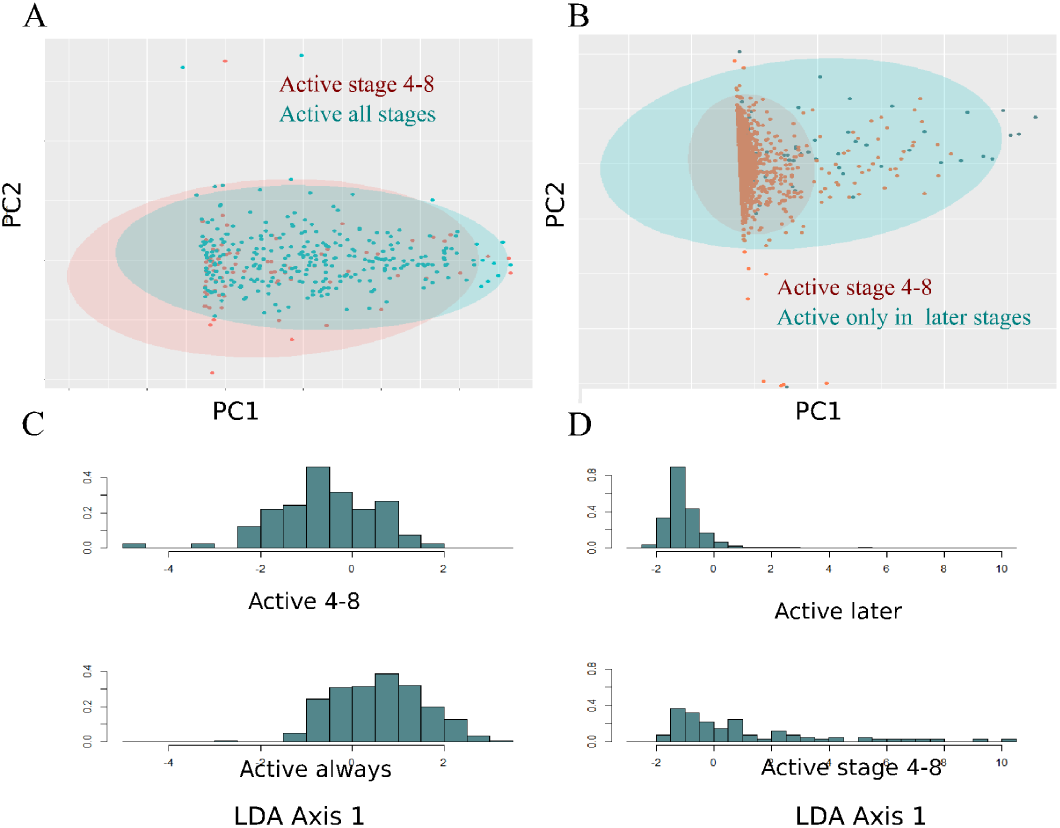
Principal component analysis, Random Forest classification, of subsets of DNA elements shows similarity of active elements. Principal components analysis shows very little separation based on the first two principal components of DNA elements that are active or inactive at specific developmental times. (a) This is true when they are separated based on continuous activation (identified as ′Always′ in Fig 7) vs. stage-specific activation (identified as ′Early′ in Fig 7), or (b) early stage specific activity from activity specific to later stages (later stages combining groups ′Middle′ and ′Late′ from Fig 8). LDA also shows very little separation along LD axis 1 for either comparison (c-d)

Predicting expression patterns is a more difficult problem than predicting activity. Although Kvon et al. 2014 annotated expression patterns for all DNA elements at each stage of embryonic development, their annotations were in many cases non-exclusive; multiple labels were applied whenever applicable. For example, a DNA element with distinct A/P stripe activity might be annotated as ″Ectoderm″ and ″Mesoderm″ as well as ″AP″, because of its partial expression in all three categories. We re-annotated expression patterns based on the images available at the Fly Enhancer Resource, grouping patterns into exclusive and non-overlapping categories. We categorized 114 elements as ″Anterior″, and 78 as ″Central or Posterior″, and 22 with ″Mesodermal or Neurogenic Ectoderm″ expression patterns. Interestingly, a substantial fraction of Anterior-Posterior and Dorsal-Ventral enhancers were ubiquitously active throughout embryonic development.

Occupancy of certain transcription factors associated with Anterior vs. Central or Posterior expression patterns correlated well with the corresponding categories of DNA elements: Bicoid peak scores were higher on Anterior elements, and Caudal peak scores were higher on Central/Posterior elements (Figure 9a and 9b). Considering all 41 features, however, PCA and LDA did not show much separation between these two expression patterns (Figure 9c, d). A random forest trained on all features performed better than predictions using randomized data, but not as well as previously observed for differentiation of active vs. inactive DNA elements (Fig. 9e). Excluding Bicoid and Caudal related data had a dramatic effect on predictions. Small training sets such as those used here are generally not compatible with large numbers of predictors, therefore, we tested dimensionally reduced datasets, to see if this would improve precision. Reducing the number of predictors using principal components did not improve performance, but in some cases degraded it (Figure 9e). Because Bicoid and Caudal play such key roles in these predictions, Anterior vs. Central/Posterior expression may be predicted well if a larger data set of specifically Anterior/Central-Posterior elements were available for training.

**Figure 9.**
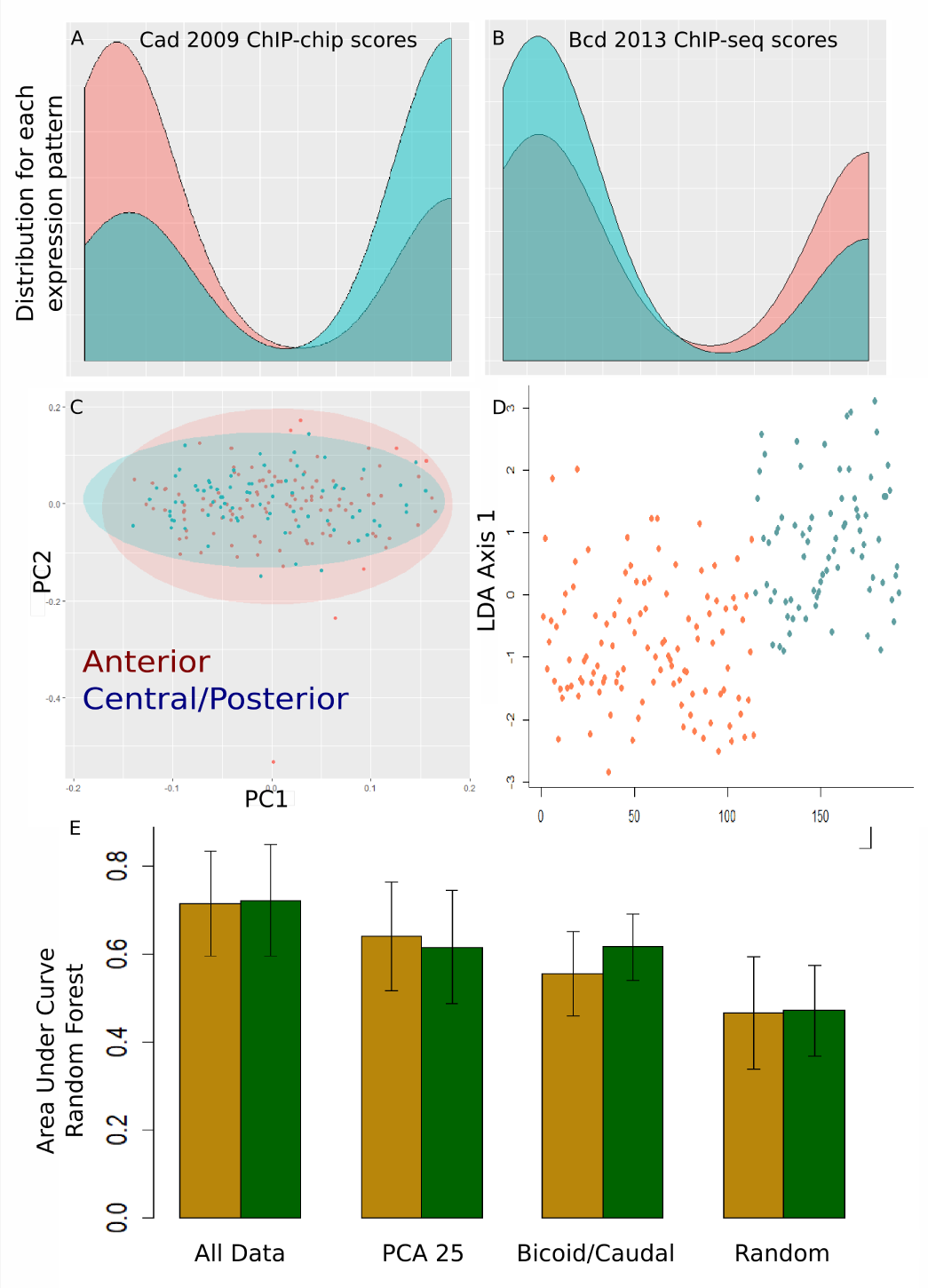
Features correlate with tissue specific expression patterns, but distinct expression patterns cannot be clearly distinguished based on principal components. Although some features clearly correlate with Anterior vs. Central/Posterior expression patterns (a,b), and these categories are largely indistinguishable when viewed by PCA (c) and show only slight separation based on LDA (d). Random forests can classify these active elements by expression pattern with greater success than when using random data, but not with sufficiently high precision or accuracy to make useful predictions. Reducing the numbers of features does little to improve accuracy of classifications, and excluding Bicoid and Caudal binding data lowered average AUC (e).

### G. Testing against validated enhancers

Enhancer classification described above depends on the extensive data from the Fly Enhancer Resource, which randomly samples 15% of the entire genome, and has limited information about which genes the putative enhancers regulate. Curated enhancers listed in the RedFly database (Gallo et al. 2011) have been experimentally associated with corresponding genes biological functions. To use these loci as independent guides for enhancer classification efforts, we tested eight different models derived from random forest analysis of balanced or unbalanced data. These models were used to predict active DNA elements within a 50kb window encompassing the eight target genes (expressed in Anterior/Posterior, Mesoderm, and Neuroectoderm patterns). Our assumption is that previously annotated regulatory regions around these genes are effective measures of true-positives, and that other regions lack enhancer activity. Thus, any additional enhancers identified within this window that do not overlap with curated enhancers are likely to be false positives.

To ensure that our random forest models evaluate all relevant regions around these genes, we needed to include additional DNA elements, as the Fly Enhancer Resource does not cover the entire genome. Therefore, we identified all putative enhancers by presence of bound transcription factors. In Figure 10, putative enhancers are defined as any locus that is binding a combination of Zelda and one or more additional transcription factors, or alternatively any locus marked by H3K4me1 as well as bound by one or more transcription factors. The ′borders′ of each putative enhancer were set to 2000 base pairs (approximately the same as the window size used in the Fly Enhancer Resources). There are 23 curated enhancers for these genes total (Table S4). However, there are an additional 31 clusters of enhancer-like binding around the protein Zelda. Almost every model trained on the Fly Enhancer Resource had high recall, and was able to identify the majority of the 23 ′true positives′ (10a). However, many conditions led to an almost equal number of ′false positives′, or a high false discovery rate (FDR) (10b). Models trained on unbalanced data were far more successful at rejecting the non-functional binding, although the balanced datasets did performed better at training and testing within the Fly Enhancer Resource. Unbalanced datasets may more accurately reflect natural conditions; the frequency of enhancer-like binding in stage 4-6 embryos appears to be far more common than actual enhancers. We performed the same test again, setting the window size around putative enhancers to 500, 1000 and 1500 base pairs, and the same trends were observed. There are far fewer clusters of binding in 2kb regions around H3K4me1 marks, which do not encompass every known enhancer. As a result, both Recall (percentage of true positives identified) and FDR were much lower.

**Figure 10.**
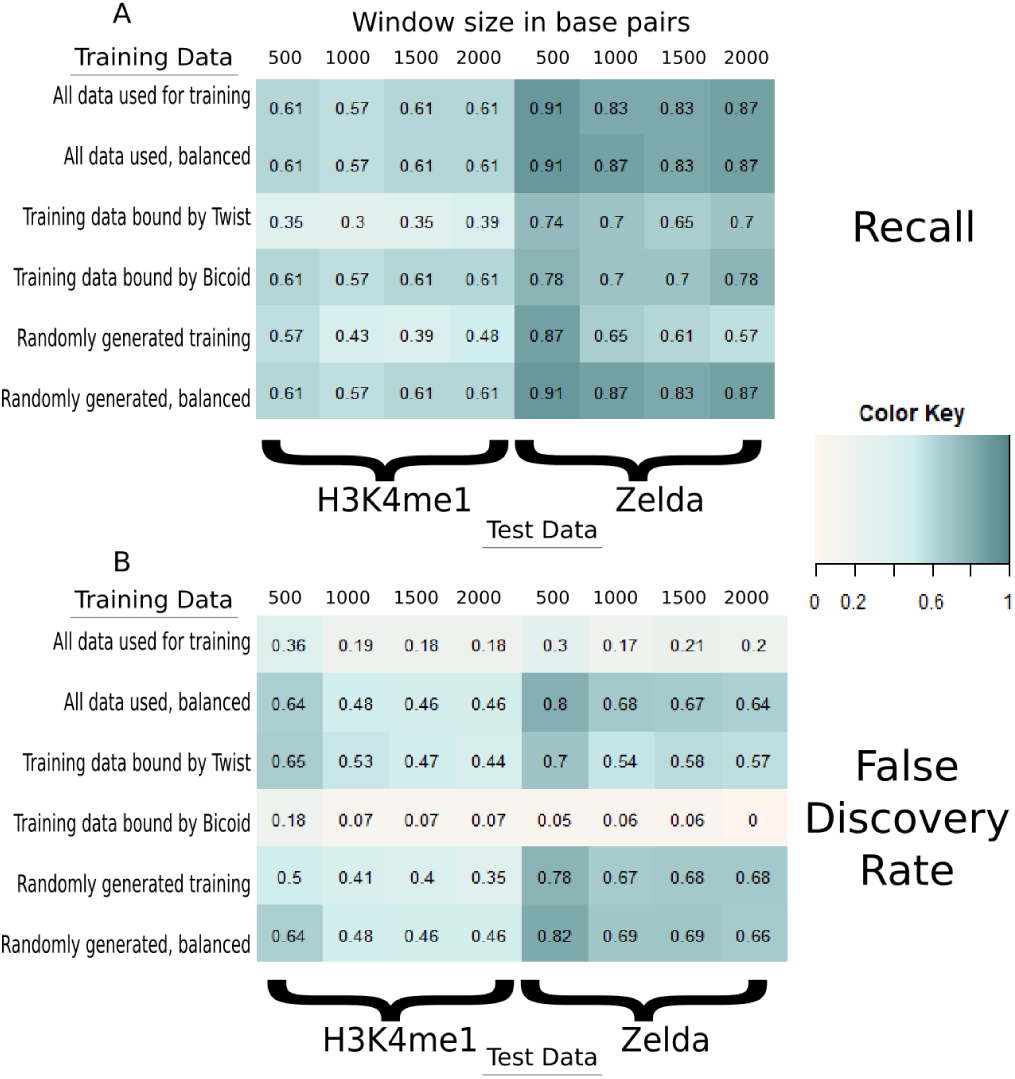
Accurate classification of active DNA elements does not consistently extend to precise or accurate identification of enhancers around well annotated genes. Recall is correlated with False Discovery Rate when identifying enhancers around well annotated genes. Clusters of transcription factors around Zelda binding OR around H3K4me1 marks, that involved two or more overlapping peaks, were used to define putative enhancers. These were defined as 500, 1000, 1500, or 2000 base pairs centered around the clustered binding. Random Forests were trained on subsets of the Fly Enhancer Resource, using all strongly active and inactive DNA elements, or only active and inactive elements that overlap with Bicoid ChIP-seq peaks, or only active and inactive elements that overlap with Twist ChIP-chip peaks (two conditions that performed very well on the Fly Enhancer Resource in Figure S7). Random forests were also trained on randomized DNA elements, where no features distinguish active and inactive elements.

## 3. DISCUSSION

More than three decades after the first description of eukaryotic enhancers, the identification and characterization of these regulatory elements is still a challenging enterprise. Initial molecular biology approaches characterized regulatory loci by ′enhancer bashing; however most regions identified as enhancers are currently products of high-throughput studies that correlate one of four marks with gene regulation: 1) protein modifications or occupancy, 2) chromatin accessibility, 3) physical association of a distal region with the basal promoter, or 4) production of enhancer RNA (eRNA). The positive correlations between these features and gene expression indicate that they can serve as good proxies for a biologically-relevant enhancer, but the extent of false positives and false negatives is generally not given enough attention. Yet it is likely that a significant portion of so-called enhancers represent biological ″chaff″, which may be the unavoidable by-product of the evolution of complex regulatory systems. For instance, the Hairy transcriptional repressor induces a large number of chromatin modifications across the genome, consistent with its ability to recruit histone deacetylases and demethylases to bound loci. Yet a minority of these sites are actually associated with changes in gene expression; the transient interactions may represent in many cases unavoidable but nugatory biochemical activity (Kok et al. 2015).

Here, to better understand our prediction of enhancers on a genome-wide scale, we took advantage of an orthogonal dataset that specifically tests the regulatory output of many genomic fragments in physiologically relevant assays. The Fly Enhancer Resource is the first publicly available database that includes a large set of reporters that are inactive at various developmental stages. The assessment of enhancer predictions against this dataset provides a rigorous, although not infallible measure of model accuracy. The data types we drew upon represent the heterogeneous quality and natures of genomic resources typically available for computational analysis of enhancers. Our conclusions about combinations of features most indicative of enhancer activity reveal the value of combinatorial types of data, as well as the serious limitations still observed.

Our results showed that active enhancers can be identified with a high degree of accuracy using genomics data. Some features (e.g. transcription factor binding in general, and pioneer transcription factor binding specifically) are far more informative than others, but combinations of less informative features are equally effective, and no one feature is essential. Classification accuracy seems to plateau after the inclusion of roughly twenty predictive features. Including more features (e.g. more ChIP data for transcription factors or histone marks) is unlikely to increase accuracy beyond where it already is. Orthogonal feature types (perhaps three dimensional data) may be necessary to break through the current upper limit on classification accuracy.

′Poised′ enhancers may be detrimental to enhancer identification; many regulatory regions assemble much of the transcriptional machinery well before the developmental time in which they are active, contributing to the false positive rate (Ghavi-Helm et al. 2014). Inclusion of predictive features that are taken from later developmental time-points may facilitate more effective pinpointing of which (if any) features can be used to identify enhancers that are active throughout development. An additional limitation of our methods is that they bundled enhancers with very different functions and characteristics into just two categories (′active′ and ′inactive′). Features that improve the accuracy of classification for elements that drive one expression pattern may be uninformative or even misleading for active elements with different functions. However, predicting expression patterns is a more difficult problem than predicting activity. A majority of the expression patterns shown in the Fly Enhancer Resource are not easily categorized; they either have multiple expression patterns evident, high variability between different embryos imaged, or very amorphous or faint distribution of protein. It is possible that these distinct tissue-specific expression patterns are more the exception than the rule at this developmental stage, and trying to group embryonic enhancers into categories based tissue specificity is not consistent with the biology of embryonic development. The distinction between non-functional and regulatory DNA, and between enhancers for one expression pattern vs. another, may be best viewed as overlapping distributions rather than distinct categories.

### A. Conclusions

Genomic features can be used to predict overall enhancer activity with a high degree of accuracy. The goals of enhancer prediction usually entail identifying as many true positives as possible, while also minimizing false-positives. Using a conservative approach when establishing the training set is key to high-precision results. A wide range of features can also lead to the best accuracy, but ChIP-seq and ChIP-chip data for a range of transcription factors and chromatin modifications seem to be far and away the most essential information. However, no single feature is a guarantee of enhancer function, and datasets measuring the same feature frequently have limited reproducibility. This has been previously observed with genome-wide studies, but it is not clear how this influences inference of function. Moreover, many features associated with a specific function (e.g. Bicoid occupancy with Anterior-Posterior patterning) can be found binding at genes associated with other regulatory networks. This may indicate functional cross-binding and integration of regulatory networks, or it could suggest promiscuous binding at regions of open chromatin. Zelda, the pioneer transcription factor, is consistently the single most important predictor of enhancer activity. However, Zelda binding is not necessary for accurate predictions. Excluding Zelda has minimal effect on overall prediction accuracy; overall accuracy appears to asymptotically level off after either a small number of highly informative features are used, or a large number of less informative features. It seems unlikely that inclusion of further ChIP-chip and ChIP-seq data would lead to meaningful improvements over the current level of performance. It also is not clear to what extent all ChIP data is serving as a proxy for DNA accessibility in these predictions, as opposed to each protein providing unique information. Although in this study motif enrichment and evolutionary conservation were not the most informative features, it is possible that more in-depth analysis in these directions, or other features that are equally orthogonal to Chromatin Immunoprecipitation data, would be necessary to reach higher levels of precision. High accuracy at predicting activity also did not translate to equally accurate identification of expression patterns. The majority of the expression patterns driven by DNA elements from the Fly Enhancer Resource were not easily categorized into clear-cut categories; the paradigm of enhancers being specific to a single gene regulatory network, and therefore to a distinct spatial expression pattern, may be biologically inaccurate.

